# Phased Diploid Genome Assembly with Single Molecule Real-Time Sequencing

**DOI:** 10.1101/056887

**Authors:** Chen-Shan Chin, Paul Peluso, Fritz J. Sedlazeck, Maria Nattestad, Gregory T. Concepcion, Alicia Clum, Christopher Dunn, Ronan O'Malley, Rosa Figueroa-Balderas, Abraham Morales-Cruz, Grant R. Cramer, Massimo Delledonne, Chongyuan Luo, Joseph R. Ecker, Dario Cantu, David R. Rank, Michael C. Schatz

## Abstract

While genome assembly projects have been successful in a number of haploid or inbred species, one of the current main challenges is assembling non-inbred or rearranged heterozygous genomes. To address this critical need, we introduce the open-source FALCON and FALCON-Unzip algorithms (https://github.com/PacificBiosciences/FALCON/) to assemble Single Molecule Real-Time (SMRT^®^) Sequencing data into highly accurate, contiguous, and correctly phased diploid genomes. We demonstrate the quality of this approach by assembling new reference sequences for three heterozygous samples, including an F1 hybrid of the model species *Arabidopsis thaliana*, the widely cultivated *V. vinifera* cv. Cabernet Sauvignon, and the coral fungus *Clavicorona pyxidata* that have challenged short-read assembly approaches. The FALCON-based assemblies were substantially more contiguous and complete than alternate short or long-read approaches. The phased diploid assembly enabled the study of haplotype structures and heterozygosities between the homologous chromosomes, including identifying widespread heterozygous structural variations within the coding sequences.

## Introduction

*De novo* genome assembly is one of the most fundamental and important computations in genome research^1–4^. It has led to the creation of very high quality reference genomes for many haploid or highly inbred species, and promoted gene discovery, comparative genomics, and several other studies^5–8^. However, despite the significance and importance of an assembly to capture the complete genetic information within an organism, most currently available genome assemblies do not capture the heterozygosity present within a diploid or polyploid species^9^. Instead, most assemblers output a “mosaic” genome sequence that can arbitrarily alternate between parental alleles^10^. Consequently, the variation between the homologous chromosomes will be lost, including allelic variations, structural variations or even entire genes present in only one of the haplotypes. Furthermore, the assemblies of heterozygous genomes tend to be more fragmented than a haploid or homozygous genome of similar size or complexity. This has greatly impeded or prevented the identification and analysis of allele specific expression, long range eQTLs, or other haplotype-specific features^11^. These challenges are becoming increasingly problematic as *de novo* sequencing projects are shifting towards more heterogeneous samples, such as outbred, wild type diploid, polyploid non-model organisms, or highly rearranged disease samples including human cancers. Beyond the intrinsic challenges of sequencing errors or repetitive elements, these samples present even more challenges for building highly contiguous assemblies, especially with short-read (<200 bp) sequencing. Consequently, the applications and utilities of such fragmented assemblies of diploid genomes have been limited^12, 13^.

While the problem of assembling diploid and polymorphic genomes is not new^14–16^, it has not been adequately solved with a generic and scalable solution. The computational methods for diploid assembly that have been proposed tend to produce highly fragmented results, often with contigs averaging just a few hundred bases to several kilobases in length ^14, 17–19^. Other approaches, such as sequencing both parents and offspring (i.e. trios) can be an effective strategy to infer haplotypes, but requires sequencing additional samples and is fundamentally limited in contiguity of the initial assemblies^20^. Sequencing individual haploid sex cells from a diploid individual is another approach that has been used^21^, but is costly and not practical for many applications. Pooled clonal fosmid sequencing^22^ produces diploid sequences but is expensive, labor intensive and the assembly contiguity is limited by the clonability of the source DNA, and the size and quality of the sequenced fosmids. Methods that rely on “synthetic long read” approaches where individual molecules are reconstructed from pooled short-read Illumina^®^ sequencing data^23, 24^ have not yielded highly contiguous sequence assemblies at the contig level in part due to limitations of resolving repeats with short reads, GC bias, and amplification bias plaguing the underlying data. Long-range scaffolding technologies (optical mapping, chromatin assays, etc.) are also no panacea for short-read diploid assembly, as they require well-assembled contig sequences (minimally contig N50 sizes 50 kbp to 100 kbp long) as a starting point, which is often out of reach for a short read assembly of heterozygous genomes.

SMRT Sequencing has now become the leading method to finish bacterial genomes and provide high contiguity assemblies for mammalian scale genomes^25–27^. The long read lengths, currently averaging ~10 kbp with some approaching 100 kbp, are fundamentally more capable of resolving repetitive elements in a genome, and should further help to resolve more complicated diploid genomes. However, currently available assemblers do not take advantage of the longer read length to resolve haplotypes. Algorithmically, though, it is possible to integrate haplotype-phasing information directly into the assembly process. In this paper, we present a new diploid-aware long-read assembler, FALCON, and an associated haplotype-resolving tool, FALCON-Unzip. They are designed to assemble haplotype contigs, “haplotigs”, representing the actual genome in its diploid state with homologous chromosomes independently represented and correctly phased^28^ (Figure 1).

**Figure 1.**
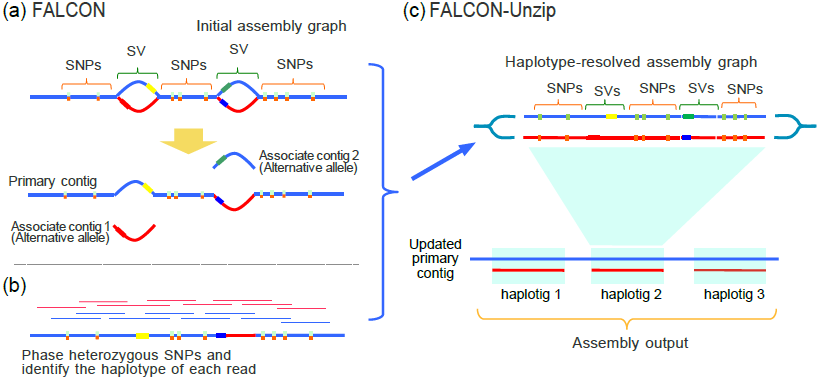
FALCON and FALCON-Unzip overview. (a) The initial assembly is computed by FALCON, which error corrects the raw reads (not shown) and then assembles using a string graph of the read overlaps. The assembled contigs are further refined by FALCON-Unzip into the final set of contigs and haplotigs. (b) Phase heterozygous SNPs and group reads by haplotype (c) The phased reads are used to open up the haplotype-fused path and generate as output a set of primary contigs and associated haplotigs.

The FALCON assembler follows the design of the previous developed Hierarchical Genome Assembly Process (HGAP)^29^, although uses greatly optimized components for each step (**Supplementary Figure 1**). FALCON begins by error-correcting PacBio raw sequence data through long-read to long-read sequence alignments and subsequently constructs a string graph^30^ of the overlapping reads. Through this process, the string graph will contain sets of “haplotype-fused” contigs as well as variant sequence “bubbles” representing (Figure 1a) major structural variations and highly divergent regions between the homologous sequences. Next, the associated tool, “FALCON-Unzip”, analyzes the “haplotype-fused” contigs and finds heterozygous variants, e.g. single nucleotide polymorphisms (SNPs) and other short variants, within the contigs (Figure 1b). It uses the phasing information of these heterozygous positions within individual reads to group the reads into different phasing blocks and haplotypes within each block. Grouped reads are subsequently re-assembled into haplotigs, and integrated with the initial “haplotype-fused” contigs in the assembly to produce the primary contigs and the haplotigs that comprise the diploid assembly (Figure 1c). The resultant haplotigs contain phased SNPs and structural variants such that a comprehensive comparison of all variants between homologous chromosomes becomes possible from the de novo assembly alone. An example of such “unzipping” process from real sequence data is shown in Figure 2.

**Figure 2.**
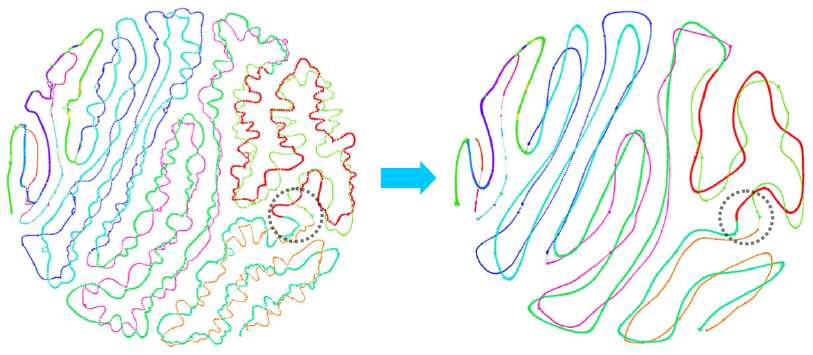
Example of Unphased and Phased Assembly Graph. (a) Initial assembly graph of a contig in the *Arabidopsis* F1 hybrid assembly. The different colors represents different haplotype blocks and phases. (b) The assembly graph after “unzipping”. Conceptually, the unzipping step identifies the heterozygous SNPs and uses them to remove overlaps between reads from different haplotypes. After removing such overlaps, nodes from the different haplotypes in the assembly graph will no longer have edges between them. This allows FALCON-Unzip to identify long haplotype specific paths and construct haplotigs of them. The dashed circle region indicates haplotype blocks that can be extended through a bubble region.

To evaluate the assembly algorithm, we sequenced two divergent inbred lines of *Arabidopsis thaliana (Arabidopsis*, henceforth), Columbia-0 (Col-0) and Cape Verde Island-0 (Cvi-0), as well as an F1 cross of the two ecotypes. The parental inbreds, and the heterozygous diploid F1 hybrid were assembled and the results were evaluated. Comparing our Col-0 assembly to the reference TAIR10 *Arabidopsis* Col-0 genome ^31^ facilitates the in depth evaluation of the assembly quality of the inbred parents. Assembling both parental genomes independently allows us to evaluate how well FALCON-Unzip can resolve the two haplotypes in the F1 genome. Once the accuracy of this process had been established, we applied the assembly algorithm on the genome of *Vitis vinifera* cv. Cabernet Sauvignon, a highly heterozygous outcrossed grape cultivar of major agricultural and economic importance. The FALCON-Unzip software suite generated a highly contiguous genome representing both sets of parental chromosomes, leading to a dramatically improved catalog of the gene content and gene duplications in the cultivar. To further demonstrate the generality of this approach, we applied FALCON-Unzip to a highly heterozygous wild-type diploid fungus, *Clavicorona pyxidata*, which has resisted previous short-read assembly approaches. For this case, we relied on orthogonal short-read and transcript sequencing data to assess the quality of the final assembly. With the transcript sequencing data, we further identified examples of differential gene expression of homologous loci that were otherwise recalcitrant to discovery by non-diploid assemblies derived from any other algorithms to date.

## Results

### Sequencing and assembly results of inbred *Arabidopsis* lines

We sequenced the inbred Col-0 and Cvi-0 genomes using 49 and 60 SMRT Cells with P4-C2 sequencing chemistry, generating 15.2 Gbp (~130x coverage) and 14.7 Gbp (~120x coverage) of raw sequence data, respectively. The raw data produced was commensurate with P4-C2 performance^32^, with average insert read length of 6.5 kbp and 6.1 kbp, and maximum read-lengths of 44,472 bp (**Supplementary Table 1**). The two inbreds were assembled independently using FALCON and both generated ~120 Mb of assembled sequence in 377 (Col-0) and 260 (Cvi-0) contigs. (Table 1). The contig N50 sizes of the assemblies were 7.4 Mb (Col-0) and 6.0 Mb (Cvi-0), about 10 to 100 times more contiguous than other recently published *Arabidopsis* assemblies produced from short-read shotgun sequencing^33^. Notably, the assembly contiguity approached that of the highly curated TAIR10 assembly of Col-0 (10.9 Mbp contig N50) that had been assembled with costly Sanger-based Bacterial Artificial Chromosome (BAC) sequencing, manual finishing, and a BAC clone tiling map^31^. Indeed, the largest FALCON contigs spanned the length of entire chromosome arms (e.g. Figure 3, **Supplementary Figure 2**), creating a new high quality reference for Cvi-0.

**Figure 3.**
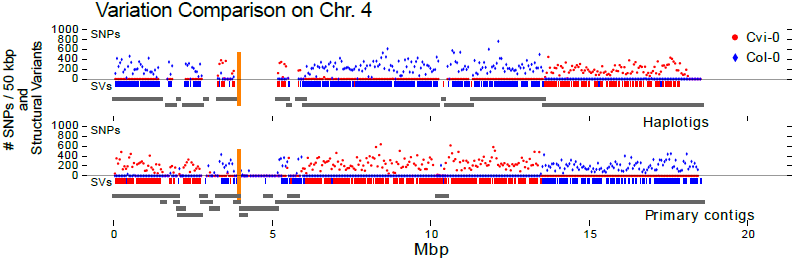
SNP density and Structural Variations in the FALCON-Unzip F1 *Arabidopsis* assembly. The plot shows the primary contigs and haplotigs aligned to chromosome 4 of the TAIR reference assembly as grey line segments. Blue and Red colored dots show the number of Col-0 and Cvi-0 specific SNPs, respectively, per 50 kbp region of the assembled contig. The vertical orange lines indicate the centromere locations. The short vertical tick marks above the grey lines indicate the structural variations against Col-0 (blue) and Cvi-0 (red).

**Table 1.**
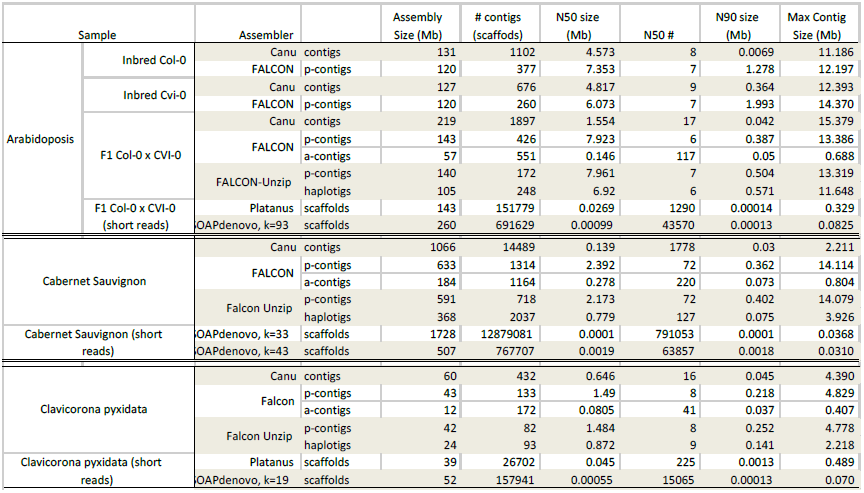
Assembly Results.

To assess the accuracy of the two inbred assemblies, the Col-0 assembly was compared with the TAIR10 assembly. We estimated the nucleotide sequence accuracy to be greater than 99.98% (**Supplementary Table 2**) between the TAIR10 reference and our Col-0 *de novo* assembly using the whole genome alignment algorithm nucmer (MUMmer3 package^34^). We applied BUSCO^35^ to evaluate the assembly completeness by identifying a set of highly conserved plant orthologs in the assembly (**Supplementary Table 3**). When BUSCO was applied to the TAIR10 reference, we found 915 (95.7%) completed orthologs out of the total set of 956. We found nearly the same numbers of orthologs in the FALCON assemblies: 914 (95.6%) in the Col-0 assembly and 906 (94.8%) in the Cvi-0 assembly.

The Col-0 and Cvi-0 genome assemblies were further compared to each other to explore the genomic variation between the two inbred lines (Table 2). The average SNP rate between the two genomes computed by nucmer was about 1 SNP every 200bp on average, over 5x fold larger than within the human population but a modest level for plant genomes. Applying the Assemblytics tool^36^ to identify structural variations between the assemblies, we found a total of ~4.7Mbp sequences affected from 1,051 structural variations greater than 50 bp (Table 2).

**Table 2.**
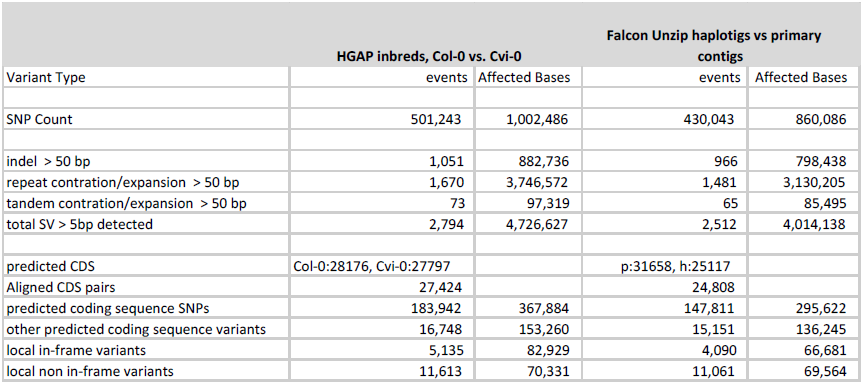
*Arabidopsis* genome assembly comparisons

| Variant Type | HGAP inbreds, Col-0 vs. Cvi-0 |  | Falcon Unzip haplotigs vs primary contigs |  |
| --- | --- | --- | --- | --- |
|  | events | Affected Bases | events | Affected Bases |
| SNP Count | 501,243 | 1,002,486 | 430,043 | 860,086 |
| indel > 50 bp | 1,051 | 882,736 | 966 | 798,438 |
| repeat contraction/expansion > 50 bp | 1,670 | 3,746,572 | 1,481 | 3,130,205 |
| tandem contraction/expansion > 50 bp | 73 | 97,319 | 65 | 85,495 |
| total SV > 5bp detected | 2,794 | 4,726,627 | 2,512 | 4,014,138 |
| predicted CDS | Col-0:28176, Cvi-0:27797 |  | p:31658, h:25117 |  |
| Aligned CDS pairs | 27,424 |  | 24,808 |  |
| predicted coding sequence SNPs | 183,942 | 367,884 | 147,811 | 295,622 |
| other predicted coding sequence variants | 16,748 | 153,260 | 15,151 | 136,245 |
| local in-frame variants | 5,135 | 82,929 | 4,090 | 66,681 |
| local non in-frame variants | 11,613 | 70,331 | 11,061 | 69,564 |

### Sequencing and assembly results of the F1 progeny of *Arabidopsis* Col-0 x Cvi-0

To assess the challenge of assembling a heterozygous genome we assembled both short and long-read sequencing data of the F1 progeny with four leading assembly algorithms (Table 1). Canu (https://github.com/marbl/canu) is an updated, although non-diploid, genome assembler based on MHAP overlapper and Celera^®^ Assembler^26^, and was used to assemble 18.5 Gbp of long-read sequence data (~ 140X haploid size from 29 SMRT Cells, length distribution shown in **Supplementary Figure 3**) from the Col-0 x Cvi-0 F1 hybrid sample. The total size of the assembly was 219 Mb, slightly smaller than the expected diploid size of 238 Mb. More significantly, the high level of polymorphisms between the two strains caused the assembly to be fragmented at regions of heterozygosity where the algorithm could not determine which of two parental assembly paths is correct. Consequently, the contiguity of the F1 assembly was substantially worse (~3 fold less) than the Canu assembly of either inbred parents alone (Table 1).

We also generated a short-read dataset (coverage = 60X, paired-end read, length = 250 bp, fragment length = 450bp) to test the short-read assemblers SOAPdenovo^37^ and Platanus^38^. The former is a widely used general-purpose short-read genome assembler, and the latter was specifically designed to assemble heterogeneous diploid genomes from short-read sequencing. The results for both assemblers were significantly less contiguous compared to Canu: SOAPdenovo assembled a total of 260 Mbp with a N50 size of 990 bp even after fc-mer optimization and error correction (**Supplementary Figure 4a and 4b**). Contigs assembled using Platanus were only marginally improved, with an N50 size of 26.9 kbp and a total assembly size of 143 Mbp, which is only slightly larger than the haploid genome size.

Unlike most genome assemblers that only generate a single set of contigs as the main assembly results, the FALCON assembler generates primary contigs (p-contigs) and associated or alternative contigs (a-contig) that comprise the genome region typified by structural variations from the primary contigs (see Methods and **Supplementary Material**). Interestingly, when the *Arabidopsis* Col-0 x Cvi-0 F1 long reads were assembled with FALCON, the assembly contiguity (p-contig N50 = 7.92 Mbp) was better than those of the *Arabidopsis* inbred-line assemblies, likely due to the longer reads used in the F1 assembly. The a-contigs, representing local alternative sequences, spanned a total of 57 Mbp (~40% of the primary contigs) with a N50 size of 146 kbp. Thus the FALCON assembler alone produced 200 Mbp (84%) of the estimated 238 Mbp diploid genome. After the initial assembly, the FALCON-Unzip algorithm utilizes the heterozygosity information within the initial primary contigs for haplotype phasing (Figure 1b, **Supplementary Material**). With the phasing information from the raw reads, it generates a subsequent set of primary contigs (p-contigs) and the final haplotig set (h-contigs) that represents more contiguous haplotype specific sequence information than the a-contigs (Figure 1c and Figure 2). After such “unzipping” process, the total size of the primary contigs was 140 Mbp (N50 = 6.92 Mbp) and the total size of the haplotigs was 105 Mbp with an N50 length of 6.92 Mbp. FALCON-Unzip nearly completely restored the contiguity that was present in the assemblies of the individual inbred parental genomes, but as a phased diploid genome.

### *Arabidopsis* Col-0 x Cvi-0 F1 haplotig phasing quality analysis

We aligned the primary contigs and the associated haplotigs to the two parental inbred assemblies to evaluate the accuracy of haplotype separations. Ideally, each haplotig should be identical to one of the parental haplotypes and show variations against the other. We observe that most of the haplotigs only show SNPs or structural variations to one of the parental genomes (Figure 3). We further analyzed the accuracy by counting the number of SNP calls against the inbred parental assemblies in each haplotig (**Supplementary Table 4**). We calculated the minority SNP percentage defined as the ratio of minimum of the two counts to the sum of the two counts. If the minority SNP percentage is small, it means there could be a small number of (1) local phasing errors, (2) incorrect SNP calls, and/or (3) assembly base errors but no major haplotype switching errors. Within the largest haplotigs that span 50% of the genome (e.g. the 6 haplotigs longer than or equal to the N50 length), the minority SNP percentages are all lower than 0.2%. Namely, there are no significant segmental switching errors inside those largest haplotigs. If, however, a phasing error occurred over multiple or large segments of a haplotig, we would expect a higher percentage of minority SNPs. Only 9 haplotigs (~2.5% of all haplotig bases) show a phasing error with a minority SNP percentage over 10%. We determined these types of assembly errors are mostly caused by repetitive regions or regions with low heterozygosity.

By design, primary contigs are only locally phased within the homologous region of each haplotig but some regions in the primary contigs have no corresponding haplotig due to low heterozygosity. FALCON-Unzip generates the primary contig to maintain the continuity through those regions but it does not maintain the haplotype phases across them. In this study, we observed that about 18.9 Mbp of separate primary contigs were actually syntenic to each other and mapped to the same region along the TAIR10 reference. This can be explained by the variations between the parental haplotypes being so extensive such that the sub-graphs of the two haplotypes are no longer associated. Due to this, the total primary assembly size may be modestly larger than the haploid genome size and the total haplotig size might be smaller than the haploid genome size.

### *Arabidopsis* Col-0 x Cvi-0 F1 base and coding sequence prediction quality analysis

In the F1 FALCON-Unzip assembly results, we estimated the overall base-to-base concordant rate at about 99.99% (QV40 in Phred scale). The insertion and deletion (indel) concordances to the parental lines were lower (about QV40) than the SNP concordance rate (about QV50) (**Supplementary Table 5**). Examining the sequence alignments around the discordant sites in detail, we found that most indel discordant sites were within long homopolymers or in simple tandem duplications. There were 68,036 A or T homopolymer blocks at least 10bp long from both haplotigs and primary contigs in our F1 assembly, and 34,483 such blocks in the TAIR10 assembly up to 48bp long (**Supplementary Figure 5**). While the SMRT Sequencing can processively read through very long, e.g. greater than 20bp, 100% A/T homopolymers, the exact length of the longest homopolymer regions may not be correct. We caution, however, that even Sanger sequencing of very long homopolymers is unreliable, and some of the discordance could arrise from errors in the reference or reflect true polymorphisms existing between the samples^39^.

To assess the quality of the assembly of the gene space, and the impact of the residual homopolymer indels on gene prediction, we first applied the gene prediction tool AUGUSTUS^40^ on TAIR10, our FALCON Cvi-0 and Col-0 assemblies, and our Cvi-0 x Col-0 F1 FALCON/FALCON-Unzip assembly and compared the predicted coding sequences (CDS) (**Supplementary Table 6**). Complete predicted CDS from the inbred and F1 assemblies aligned completely to 95-97% of the 27,954 CDS of TAIR10 without indels. To avoid potential bias due to *ab initio* prediction, we performed an additional analysis by directly aligning the manually curated TAIR10 CDS to the entire assembly using the STAR aligner^41^. Overall, 96% of the 35,386 TAIR10 CDS were fully aligned without indels or truncations to the F1 assembly contigs. Another 1.04% CDS were successfully aligned allowing for only one base insertion or deletion: 0.18% CDS had a 1 base insertion and 0.86% CDS had a 1 base deletion.

We also applied BUSCO to further assess the completeness of the assembled gene space. BUSCO found 906 single copy highly conserved orthologs in the F1 assembly, the same number that could be found in the Cvi-0 and only slightly fewer than found in Col-0 (see above). The small reduction was most often because the current phasing algorithm is only applied to primary contigs longer than 20 kbp, and initial primary contigs shorter than 20 kbp are filtered out. Furthermore, since the assembly contains both haplotypes (i.e. primary contigs and haplotigs), the vast majority of BUSCO genes (877) were found to be duplicated in the assemblies showing that the majority of the gene space was phased by the diploid assembly.

### Analysis of the variations between haplotypes in *Arabidopsis* Col-0 x Cvi-0 F1 genome

With both haplotype sequences robustly assembled, we could analyze the differences between the homologous chromosomes using nucmer and Assemblytics (Table 2). When we compared the haplotigs to the primary contigs in the F1 assembly, we identified 430,043 SNPs, compared to 501,243 found by aligning the Col-0 and Cvi-0 inbred assemblies. Using Assemblytics, we identified 966 SV events (>50bp indels or tandem repeat contractions and expansions) between the haplotigs and primary contigs, compared to 1,051 between the Cvi-0 and Col-0 assemblies. Thus, FALCON-Unzip phased 85.7% of all SNPs and 91.9% of all SVs directly from the shotgun sequence assembly. Interestingly, nearly one third of the 31,679 Augustus predicted coding regions intersected structural variants at least 50bp in size identified on the primary contigs, which may have important effect on gene expression regulation and/or functionality.

We estimated the amount of variation affecting coding sequences by comparing the predicted CDS (Table 2) between the Col-0 and Cvi-0 inbred lines. We found about 184,000 (0.45%) SNPs within the 40.7Mbp predicted CDS of the inbred assemblies, compared to 148,000 (0.41%) SNPs within the 36 Mbp CDS between the haplotigs and the primary contigs in the FALCON-Unzip assembly of the F1 (Table 2). The number of heterozygous SNPs and SVs present in the F1 assembly is marginally lower than those from the comparison between Col-0 and Cvi-0, mostly because the collection of the haplotigs does not fully represent the full haploid chromosome set. In particular, the number of variants between the haplotigs and the primary contigs is consistent with the total haplotig size (105 Mbp) that is about 87% of the estimated genome size.

### Comparison of the long-read and short-read assemblies of the *Arabidopsis* Col-0 x Cvi-0 F1 genome

Our final analysis was to compare the short and long read assemblies to the TAIR10 genome to assess the quality differences between the assemblies and the ability to identify variants. We used Assemblytics to call insertion and deletion variants from the three *Arabidopsis* F1 assemblies with FALCON-Unzip, Platanus, and SOAPdenovo to the TAIR10 reference. We aligned the contig sequences for the short-read assemblies since aligning the scaffolds may introduce artificial variants due sequence gaps marked with Ns. Assemblytics identified a total of 215,801 variants from the FALCON-Unzip assembly, of which 3,847 were structural variants larger than 50bp (**Supplementary Figure 6a/b**, **Supplementary Table 7**). In contrast, Assemblytics detected 85,899 variants (1,128 sites >50bp) in the Platanus assembly and only 2,414 variants (10 sites >50bp) from the SOAPdenovo assembly. The variants from the FALCON-Unzip assembly captured 89% of the Platanus variants and 90% of the SOAP variants at a stringent requirement of the exact same variant type, size, and genomic location. However, the Platanus and SOAP assemblies captured only 37% and 1% of the FALCON-Unzip variants, respectively. The contiguity of the assemblies greatly affects the numbers and sizes of variants that can be called, and since differing haplotypes can result in mis-assemblies, constructing proper haplotypes can be an important factor in accurate variant-calling from an assembly.

### *V. vinifera cv*. Cabernet Sauvignon sequencing and diploid assembly results

After establishing the accuracy of the assembly pipeline with *Arabidopsis*, we assessed the performance of FALCON-Unzip on the genome of *V. vinifera* cv. Cabernet Sauvignon, a highly heterozygous outcrossed grape cultivar. Cabernet Sauvignon is an F1 of two very distinct cultivars, Cabernet Franc and Sauvignon Blanc^42^ and one of the world’s most widely cultivated red wine grape varieties. We sequenced 74 SMRT Cells using the PacBio^®^ P6-C4 chemistry yielding 73.7 Gbp of sequence, equivalent to ~140X of the haploid genome, with average read length of 10.7 kbp (**Supplementary Table 1**). Reads were assembled using Canu, FALCON, and FALCON-Unzip (Table 1). FALCON-Unzip yielded the most contiguous assembly with a N50 size of 2.17 Mbp and generated a total of 368 Mbp of associated haplotigs with N50 of 779 kbp. Both primary and associated contigs displayed overall high macro-synteny with the current *V. vinifera* genome reference (PN40024^43^; **Supplementary Figures 7 a/b**). The total primary assembly size (590 Mbp) was larger than the estimated genome size of *V. vinifera* (~500Mbp^43^). This suggests that in some cases FALCON-Unzip underestimated the alternative haplotype sequence, because of incomplete bubble structure in the assembly graph. An analysis of synteny between primary contigs to determine the extent of inclusion of redundant regions in the primary assembly identified a total of 25 Mbp of syntenic blocks (see **Supplementary Material**).

The more complicated repeat and diploid structure^43^ posed more of a challenge for the typical non-diploid aware long-read and short-read whole genome shotgun assembly approaches. We tested the Canu assembler on the same PacBio reads, and SOAPdenovo and Platanus on a short-read dataset consisting of 45x coverage of the haploid genome size in paired-end 100bp reads. Similar to *Arabidopsis*, Canu generated an assembly of 1,006 Mbp which is roughly twice of the haploid genome size with a significantly smaller N50 size of 139 kbp. With the short-read data, the assembly N50 and total assembly size of the SOAPdenovo assembly were sensitive to the choice of fc-mer size for constructing the de Bruijn graph (**Supplementary Figure 4a/b**). Even with optimized fc-mer sizes (33-43), the scaffold N50 sizes were smaller than 2 kbp and the contig N50 sizes were less than 1 kbp. The Platanus results were unacceptably incomplete, with less than 1% of the expected genome size reported, most likely due to the limited available coverage. However, even under idealized conditions with higher coverage levels (1,577 million reads) and multiple libraries, other published assemblies of different grape cultivars report contig N50 sizes of at most 41 kbp using Platanus^44^.

To assess completeness of the assemblies we applied the BUSCO pipeline as well as aligned the 29,971 mRNA sequences annotated from the current *V. vinifera* genome reference PN40024. Both approaches highlighted the completeness of the gene space in the FALCON-Unzip assembly, similar to what we observed for *Arabidopsis* (**Supplementary Table 3, 6 and 8**). We extended this analysis to determine the representation of the gene space in the associated assembly. Overall 80% of the 766 duplicated BUSCO proteins were found with copies in both primary and associated haplotigs, while 16,981 complete genes from PN40024 were in common between primary and associated haplotigs. In contrast, less than 15% of the 956 BUSCO proteins were found within the most contiguous short-read assemblies suggesting that these assemblies are not only highly fragmented, but also egregiously incomplete (**Supplementary Table 3**).

### *Clavicorona pyxidata* sequencing and assembly results

To demonstrate the generality of the FALCON-Unzip approach to wild type heterozygous genomes, we assembled *C. pyxidata*, a common coral fungus that grows on hardwoods across North America (haploid size ~42 Mbp). A total of 4.0 Gbp (~100X haploid size) of data from 6 SMRT Cells was used to produce a diploid assembly of *C. pyxidata* (see **Supplementary Figure 3** for the read length distribution). Similar to the two plant genomes, a long-read assembly was performed with FALCON/FALCON-Unzip and Canu as well as two short-reads assemblies with Platanus and SOAPdenovo. The overall assembly results were similar to those observed with the plant assemblies (Table 1): FALCON-Unzip produced the most contiguous assembly, followed by Canu (~2-fold less contiguous), and then followed distantly by the short-read assemblies (30 to >100 fold less contiguous). Interestingly, about half of the genome was found to have significantly reduced rates of heterozygosity between the homologous chromosomes, suggesting naturally occurring inbreeding events or other selective pressure to maintain homozygosity through part of the genome. Thus, the haplotigs spanned about 50% of the genome.

Unlike the case with the *Arabidopsis* and *V. vinifera* assemblies, there is no highly contiguous and curated reference sequence to evaluate the completeness or base level accuracy by assembly-assembly alignment for *C. pyxidata. In lieu* of a reference, we evaluated the completeness of the gene space using the BUSCO and genomic sequencing data from an orthogonal sequencing platform (SRA accession: SRR1800147, 86X, 150 bp reads) to evaluate the assembly accuracy (**Supplementary Table 3 and 9**). The scaffold N50 (45 kbp) of the Platanus assembly was 30 times smaller than FALCON-Unzip’s primary contigs’ N50 (1.48 Mbp), but the Platanus assembly did contain approximately the same number of single copy eukaryote BUSCO sets as those in the FALCON-Unzip assembly (**Supplementary Table 3**). However, the Platanus assembly missed nearly all of the homologous copies in the diploid genome (29/429 eukaryotic BUSCO proteins are duplicated compared to 277/429 in the FALCON-Unzip assembly) and the regulatory context around the genes are much more limited. The best SOAPdenovo assembly (with fc-mer size 19) yields less then 3% single complete genes copies and had a shorter N50 than the Platanus assembly.

We utilized the short-read sequence for a genome-wide evaluation of the assembly and phasing accuracy even without the parental assemblies or previous assembly of closely related species. With the 150bp paired-end reads, we called phased SNPs relative to the primary contigs with FreeBayes^45^ and HapCut^46^. The FALCON-Unzip assembled contigs had 0.3% to 0.6% phase discordant events between neighboring pairs of SNPs from the short-read datasets, depending on the variant call quality filters (**Supplementary Table 9**). However, due to the insert size limit of the short-read dataset, the phasing data from the short reads only covered about 23% (9.72 Mbp) of the genome. Nearly all phased blocks, 96% to 98% depending on the variant call quality thresholds, are fully concordant with the assembly (**Supplementary Table 9**). This indicates the short range phasing is consistent between the FALCON-Unzip assembly and the short-read data, although the long reads produce much longer phased regions. Comparison of homologous alleles within the genome with public available RNA Sequencing data (SRA accession SRR1589642) identified several candidate differentially expressed alleles (**Supplementary Figure 8a-8e**).

## Discussion

The aim of many genome sequencing projects is to generate a high quality reference assembly that can serve as a foundation for various downstream analyses, e.g. gene finding, variant identification, or comparative & functional assays. While successful in a number of haploid or inbred species, one of the current main challenges for the genomics community is developing genome assemblies for noninbred heterozygous genomes, especially since these represent the vast majority of samples to be sequenced for biomedical, agricultural, or evolutionary studies. For heterozygous diploid genomes, we demonstrated FALCON and FALCON-Unzip can assemble PacBio SMRT Sequencing data into highly accurate, contiguous, and correctly phased primary contigs and associated haplotigs. Such haplotype specific assemblies present a true representation of the genome and empower study of haplotype structures and heterozygosities, e.g. structural variations and SNPs, between the homologous chromosomes not normally possible from other assemblers.

In all three genomes studied here, the FALCON/FALCON-Unzip assembly was significantly more contiguous (2 to 3 fold) than alternative long read assemblers of the same data, and overwhelmingly better (30 to >100 fold) than state-of-the-art short read assemblies. In the *Arabidopsis* F1-hybrid assembly, we evaluated the haplotype phasing accuracy by comparing the F1 assembly to the parental inbred genomes and determined that the haplotigs nearly perfectly matched one of their parental genomes with only a very small number of incorrectly phased alleles. While already extremely accurate, in future work, we aim to further improve the phasing accuracy by analyzing the local assembly graph to predict hard-to-resolve regions and potential errors in the assembly. We further showed that the small frequency of residual sequencing errors (<0.1%) had no substantial effect on the ability to correctly identify gene sequences. In the other two assemblies, we demonstrated greatly improved diploid representations of core genes from the FALCON/FALCON-Unzip assembly, and highly accurate phasing measured using orthogonal data.

At a fundamental level, both the raw sequencing read lengths and error rates may affect the haplotype and consensus accuracies. The genome content, especially the rate of heterozygous positions and the repetitive sequences, is also a major factor impacting the performances. Most haplotype-phasing algorithms utilize heterozygous SNPs and ignore any structural variations. In contrast, an overlap-layout-consensus genome assembly paradigm captures structural variations between haplotypes naturally as bubble structures in assembly graphs. FALCON-Unzip is designed to combine SNPs and SVs to separate haplotype information beyond what either alone provides to construct haplotype specific contigs. With long read lengths from SMRT Sequencing and increased levels of heterozygosity, this allows us to almost fully resolve both haplotype chromosomes for practically the entire Arabidopsis F1 genome with high contiguity. The other two genomes chosen for this study highlight some of the additional complexities that are possible for diploid genomes. In *V. vinifera*, we find homologous regions having very high rates of variations, likely from the out-crossing nature of the organism, while in *C. pyxidata* we discovered extended regions of unexpectedly low heterozygosity suggesting regions of increased selective pressures or complex naturally occurring inbreeding. While future increase of the read lengths will improve the separation of the haplotypes, we can already begin to utilize the assembly output to understand and represent the variations of heterozygosity within most diploid genomes across the tree of life. The assembly results presented here were solely from PacBio SMRT Sequencing, but can in principle be also improved with other types of data, especially long range scaffolding data, and extended to higher ploidy genomes in the future.

The mosaic genome sequences that are commonly assembled today do not contain all of the genetic information of the heterozygosity between haplotypes. This makes it, among other things, difficult to probe the impact of epigenetic and differential gene expression and can exacerbate “reference-bias” when remapping sequencing data^47^. With FALCON-Unzip, however, almost the entire heterozygosity information is captured in the primary contigs and haplotigs, so the question of how haplotype specific variations affect gene expression, methylation patterns, or other regulatory interactions can be examined further. We anticipate more systematic study of phased diploid references will expose the detailed cis-regulatory mechanisms of differential expression in diploid genomes to improve our general understanding of the biology beyond haploid genomes. In summary, with the advances of the SMRT Sequencing technology, new algorithm and software development, we expect that there is a wide field of new opportunities for understanding diploid and polyploid genomic diversity and its impact on genome annotation, gene regulation and evolution.

## Materials and Methods

### DNA isolation and library preparation

For the *Arabidopsis* sample preparation, to minimize chloroplast DNA contamination, nuclei were isolated from leaf tissue as previous described^48^. Genomic DNA was isolated using standard purification columns and protocols (Qiagen^®^). For grapevine DNA extraction, young leaves (~1 cm diameter) were collected from *Vitis vinifera* cv. Cabernet Sauvignon clone 08 at Foundation Plant Services (UC Davis, Davis, CA). Plant tissue (1 g) was ground to a powder in a mortar containing liquid nitrogen. Ten mL of pre-warmed (65 °C) extraction buffer (300 mM Tris-HCl pH 8.0, 25 mM EDTA pH 8.0, 2 M NaCl, 2% (w/v) soluble PVP (MW 40000), 2% CTAB, 2% 2-mercaptoethanol) was added and the suspension was homogenized by inversion and incubated (65°C) for 30 min in a water bath, mixing by inversion (every 5 min). Plant debris was removed by centrifugation (5000 rpm) for 5 min at room temperature and the supernatant was transferred into a new tube. Equal volume of chloroform:isoamyl alcohol (CIA, 24:1 v/v) was added and mixed by inversion for 5 min. Aqueous phase was segregated by 10 min centrifugation (5000 rpm) at room temperature and transferred gently into a new tube. RNase A was added to the sample (2 μg) and was incubated (37°C) for 30 min. After RNAse treatment, equal volume of CIA was added and centrifuged as above. 0.1 volume of 3 M NaOAc pH 5.2 and an equal volume of isopropanol were added for DNA precipitation, sample was mixed by inversion and then incubated (-80 °C) for 30 min. DNA was collected by centrifugation (5000 rpm) for 30 min and the pellet was washed twice with 3 mL of 70 % ethanol. After 10 min centrifugation (5000 rpm), DNA pellet was air-dried at room temperature and resuspended in 500 μl of nuclease-free water. DNA quality was evaluated by pulse-gel electrophoresis, and quantity was determined using the Qubit fluorometer.

Shearing of the DNA was performed either with G-tubes (Covaris^®^) or by passage through a small bore needle ^49^ to average size of 15 kbp to 40 kbp. Sheared DNA was enzymatically repaired and converted into SMRTbell™ libraries prepared as described by the manufacturer (Pacific Biosciences). Non SMRTbell DNA was removed by exonuclease treatment. Finally, a BluePippin™ preparative electrophoresis purification step was performed (Sage Sciences) on the library to select insert sizes ranging from 7 to 50 kbp or from 15 to 50 kbp depending on the sequencing experiment. These size-selected libraries were used in subsequent sequencing steps.

### Sequencing methods

Sequencing was performed on the PacBio RS II instrument as per the manufacturer’s recommendations. The Col-0 and Cvi-0 inbred *Arabidopsis* data sets were collected using P4-C2 chemistry with 4 hour movie lengths. The F1 Col-0 x Cvi-0 and the *C. pyxidata* and the *V. vinifera* cv Cabernet Sauvignon samples were run with P6 chemistry and 6 hour data collection movies.

### Raw long-read error correction

All raw long-read sequences were aligned to each other using “daligner^50^” executed by the main script of the FALCON assembler. The overlap data and raw subreads are then processed to generate consensus sequences. The consensus-calling algorithm (FALCON-sense) was designed to preserve the information from heterozygous single nucleotide polymorphisms (SNP) and is described in detail in the **Supplementary Material**.

### Initial “haplotype-fused” assembly with a collapsed diploid-aware contig layout

After the error correction step, FALCON identifies the overlaps between all pairs of the preassembled error corrected reads. The read overlaps were used to construct a directed (in contrast to bidirected) string graph following the Myers’ method^30^. For diploid genomes with high heterozygosity, the string graph typically contains linear chains of “bubbles” (Figure 2 and **Supplementary Figure 13**). We can decompose such linear chains into “simple” and “compound” paths where: a simple path is a path where there is no internal branching node and it also has unique source node and sink node, and a compound path is a collection of edges that represents a bubble with unique source and sink in the assembly graph. The algorithm for constructing such compound paths is described in the **Supplementary Material**. The non-branched collection of compound paths and simple paths are further combined to create unitigs. Genome repeats, sequencing errors or missing overlaps can introduce spurious unitigs. Empirically derived heuristic rules were applied to remove these artifacts and layout the primary contigs and the associated contigs. The graph reduction process is detailed in **Supplementary Figure 10**. We call the final assembly graph the “haplotype-fused assembly graph.”

### Mapping and phasing the raw reads

In the draft assembly, each contig is simply a tiling sequence from the subsequences of a set of error corrected reads. Some of the raw reads have not yet been associated with any contigs. For example, if a read is “contained” within other reads (overlaps completely to a substring of another read), it is not used in constructing the first draft of the contigs. There are two strategies for identifying the raw-read to contig associations: (1) re-map all raw-reads to the contigs and find the best alignments; or (2) trace the read overlapping information to find out where a raw-read is mostly likely to be associated. FALCON-Unzip applies strategy (2), to avoid the time penalty for the re-mapping process, as the overlap information already exists. For each raw-read, FALCON-Unzip examines all overlapping reads. If a read is uniquely associated with one contig, then the raw-read is assigned to that contig. If there are multiple contigs associated with a read, it scores the matching contigs by the overlap lengths. In this case, a read is assigned to a target contig with the highest sum of overlap lengths.

For each primary contig, we collect all raw-reads associated with the primary contig and its associated contigs. We align the raw reads to the contigs with the BLASR aligner^51^ and call heterozygous SNPs (het-SNPs) by analyzing the base frequency of the detailed sequence alignments. A simple phasing algorithm was developed to identify phased SNPs (see **Supplementary Material**). Along each contig, the algorithm assigns phasing-blocks where chained phased SNPs can be identified. Within each block, if a raw read contains a sufficient number of het-SNPs, it assigns a haplotype phase for the read unambiguously. Combined with the block and the haplotype phase information, it assigns a “block-phase” tag for each phased read in each phasing block. Some reads might not have enough phasing information. For example, if there are not enough het-SNP sites covered by a read, it assigns a special “un-phased tag” for each un-phased read.

### Overview of the algorithm constructing haplotype specific contigs

The algorithm to construct the haplotype specific contigs (haplotigs) is summarized in Figure 1 and **Supplementary Figure 12 and 13**. Briefly, for each contig *c*, it constructs a haplotype-specific assembly graph from all reads that mapped to it, denoted as *H_c_*, by ignoring the overlaps between any two reads from the same block but different phases. It then combines this graph *H_c_* to the fused assembly sub-graph 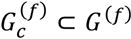 that contains the paths of contig *c* to construct a complete contig subgraph 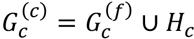 Unlike the initial subgraph 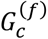, where some reads are masked out by reads fc) from different phases, the complete contig sub-graph 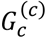 rescues such masked-out reads and have complete read representation from both haplotypes.

In the fused assembly graph 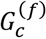, there is a path that is corresponding to the original contig *c* starting from node *s* to node *t*. It is desirable to generate a new locally phased contig that also starts from the same node *s* and ends at the same node t as new primary contig *p_c_*. While such primary contig *p_c_* may not be fully phased end-to-end, the collection of *p_c_* of all contig *c* can serve as a haploid assembly representation with annotated locally phased regions. And, the variations between the two haplotypes can be identified by aligning other haplotigs to the primary contigs. Once *p_c_* is identified, the corresponding edges of *p_c_* in 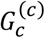 are removed. It also removes all other edges connecting different phases of the same block. Namely, it constructs a subgraph 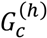 of 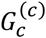 by removing edges which are already in *p_c_* or connect distinctly phased nodes. We identify all linear paths within 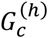 as the haplotigs *h*_*c,i*=1‥*n*_, where *n* is the total number of haplotigs associated with the primary contig. Some of the haplotigs might be caused by missing overlaps or sequence errors. The haplotig sequences are aligned to the primary contig. If the alignment identity is high and no phased-reads are associated with the haplotig, the haplotig will be marked as duplicated and removed. Note that a haplotig may contain multiple haplotype-phased blocks. For example, haplotype-specific structural variation may affect the initial mapping such that the phasing algorithm cannot connect two neighboring blocks. However, reads from different phasing blocks might be uniquely overlapped if the structural variations between the haplotypes are distinguishable. Such haplotype-specific overlaps can connect broken haplotype-phased blocks into to larger haplotigs.

### Polishing partially phased primary contigs and their associated haplotigs

Conceptually, FALCON-Unzip generates one new primary contig *p_c_* and *n* haplotigs *h*_*c,i*=1‥*n*_ from the original assembly graph 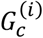 of the contig *c*. It uses the phasing information to decide whether a phased read belongs to the primary contig *p_c_* or one of the haplotigs *h*_*c,i*=1‥*n*_. Each un-phased read may also contain structural level variations that are the same as in a particular haplotig. In such case, by examining the overlaps between the read to those in the haplotigs, it can find the best hit from the unphased read to one haplotig. In the end, each raw-read will be augmented with the information which haplotig or primary contig it belongs to and will be mapped accordingly. This ensures that the haplotig consensus is generated from the appropriate reads belonging to the correct haplotype. Finally, it uses the Quiver algorithm^29^ to remove residual errors in the haplotig consensus from the haplotype specific alignments.

FALCON-Unzip outputs a set of partially phased primary contigs (p-contigs) and the associated haplotigs (h-contigs) for each primary contig. The phased regions in the primary contig can be identified by simply aligning the associated haplotigs to the primary contig or directly examine the assembly graph identifying the anchoring nodes from the haplotigs to the primary contig.

## Software Availability

FALCON and FALCON-unzip are written in C and Python. FALCON and its dependences are hosted open-source on GitHub^®^ (https://github.com/PacificBiosciences/falcon). FALCON-Unzip is also hosted open-source on GitHub^®^(https://github.com/PacificBiosciences/FALCONunzip). The specific git repositories of the various modules used for generating the assemblies presented in this paper are listed in the **supplementary material**. We have also prepared an Amazon Web Services EBS volume that contains all of the preconfigured software and example *C. pyxidata* dataset (See **Supplementary Materials** for a walkthrough).

## Assembly Files

The assemblies can be downloaded from https://downloads.pacbcloud.com/public/dataset/PhasedDiploidAsmPaperData/FUNZIP-PhasedDiploidAssemblies.tgz. The GenBank accession numbers are in progress.

## Data Accession

*Arabidopsis* data: PRJNA314706

*V. vinifera* cv. Cabernet Sauvignon: PRJNA316730

*Clavicorona pyxidata:* Upload to SRA in progress.

## Acknowledgements

The sequencing of the Cabernet Sauvignon genome was partially supported by a gift of the J. Lohr Vineyards to DC. We also like to thank Felipe Simao Neto providing early release BUSCO plant data set. *Clavicorona pyxidata* DNA was provided by Laszlo Nagy. We thank Joseph D. Puglisi, Florian Jupe, and Alex Copeland for reading and critique of the manuscript. The project was supported in part by National Institutes of Health award (R01-HG006677) and by National Science Foundation awards (DBI-1350041 and IOS-1237880 to MCS; MCB 0929402 and MCB 1122246 to J.R.E)/ J.R.E is an investigator of the Howard Hughes Medical Institute and Gordon and Betty Moore Foundation (GBMF 3034).

## Supplementary Table Captions

Supplementary Table 1. Raw sequence data read length statistics

Supplementary Table 2. Concordance of *Arabidopsis* TAIR10 with Falcon assembly of *Arabidopsis* Col-0

Supplementary Table 3. BUSCO results for all assemblies

Supplementary Table 4. Haplotig SNP rate against two parental inbred lines

Supplementary Table 5. Haplotig concordance against two parental inbred lines

Supplementary Table 6. *Arabidopsis* AUGUSTUS CDS prediction results

Supplementary Table 7. Comparison of structural variation calls of F1 long and short-read assemblies to TAIR10

Supplementary Table 8. *V. vinifera* CDS alignment summary

Supplementary Table 9. FALCON-Unzip phase concordance with short-read dataset

## References

1. Goffeau, A. et al. Life with 6000 genes. Science 274, 546–567 (1996).

2. Adams, M.D. et al. The genome sequence of Drosophila melanogaster. Science 287, 2185–2195 (2000).

3. Myers, E.W. et al. A whole-genome assembly of Drosophila. Science 287, 2196–2204 (2000).

4. Bonfield, J.K., Smith, K.F. & Staden, R. A new DNA sequence assembly program. Nucleic acids research 23, 4992–4999 (1995).

5. Stamatoyannopoulos, J.A., Guigó Serra, R., Djebali, S., Lagarde, J. & Adams, L.B. An encyclopedia of mouse DNA elements (Mouse ENCODE). (2012).

6. Consortium, E.P. An integrated encyclopedia of DNA elements in the human genome. Nature 489, 57–74 (2012).

7. Celniker, S.E. et al. Unlocking the secrets of the genome. Nature 459, 927–930 (2009).

8. Consortium, G.P. A global reference for human genetic variation. Nature 526, 68–74 (2015).

9. Earl, D. et al. Assemblathon 1: a competitive assessment of de novo short read assembly methods. Genome research 21, 2224–2241 (2011).

10. Church, D.M. et al. Extending reference assembly models. Genome biology 16, 13 (2015).

11. Tewhey, R., Bansal, V., Torkamani, A., Topol, E.J. & Schork, N.J. The importance of phase information for human genomics. Nat Rev Genet 12, 215–223 (2011).

12. Henson, J., Tischler, G. & Ning, Z. Next-generation sequencing and large genome assemblies. Pharmacogenomics 13, 901–915 (2012).

13. Alkan, C., Sajjadian, S. & Eichler, E.E. Limitations of next-generation genome sequence assembly. Nature methods 8, 61–65 (2011).

14. Vinson, J.P. et al. Assembly of polymorphic genomes: algorithms and application to Ciona savignyi. Genome Res 15, 1127–1135 (2005).

15. Levy, S. et al. The diploid genome sequence of an individual human. PLoS Biol 5, e254 (2007).

16. Jones, T. et al. The diploid genome sequence of Candida albicans. Proceedings of the National Academy of Sciences of the United States of America 101, 7329–7334 (2004).

17. Donmez, N. & Brudno, M. in Proceedings of the 15th Annual international conference on Research in computational molecular biology 38–52 (Springer-Verlag, Vancouver, BC, Canada; 2011).

18. Iqbal, Z., Caccamo, M., Turner, I., Flicek, P. & McVean, G. De novo assembly and genotyping of variants using colored de Bruijn graphs. Nature genetics 44, 226–232 (2012).

19. Kajitani, R. et al. Efficient de novo assembly of highly heterozygous genomes from whole-genome shotgun short reads. Genome research 24, 1384–1395 (2014).

20. Roach, J. et al. Chromosomal haplotypes by genetic phasing of human families. Am J Hum Genet 89, 382–397 (2011).

21. Kirkness, E. et al. Sequencing of isolated sperm cells for direct haplotyping of a human genome. Genome Res 23, 826–832 (2013).

22. Kitzman, J. et al. Haplotype-resolved genome sequencing of a Gujarati Indian individual. Nat Biotechnol 29, 59–63 (2011).

23. McCoy, R.C. et al. Illumina TruSeq synthetic long-reads empower de novo assembly and resolve complex, highly-repetitive transposable elements. (2014).

24. Mostovoy, Y. et al. A hybrid approach for de novo human genome sequence assembly and phasing. Nature Methods (2016).

25. Koren, S. & Phillippy, A.M. One chromosome, one contig: complete microbial genomes from long-read sequencing and assembly. Current opinion in microbiology 23, 110–120 (2015).

26. Berlin, K. et al. Assembling large genomes with single-molecule sequencing and locality-sensitive hashing. Nature biotechnology (2015).

27. Gordon, D. et al. Long-read sequence assembly of the gorilla genome. Science 352, aae0344 (2016).

28. Makoff, A.J. & Flomen, R.H. Detailed analysis of 15q11-q14 sequence corrects errors and gaps in the public access sequence to fully reveal large segmental duplications at breakpoints for Prader-Willi, Angelman, and inv dup (15) syndromes. Genome biology 8, R114 (2007).

29. Chin, C.-S. et al. Nonhybrid, finished microbial genome assemblies from long-read SMRT sequencing data. Nature methods 10, 563–569 (2013).

30. Myers, E.W. The fragment assembly string graph. Bioinformatics 21, ii79–ii85 (2005).

31. Initiative, A.G. Analysis of the genome sequence of the flowering plant Arabidopsis thaliana. nature 408, 796 (2000).

32. Kim, K.E. et al. Long-read, whole-genome shotgun sequence data for five model organisms. Scientific data 1 (2014).

33. Gan, X. et al. Multiple reference genomes and transcriptomes for Arabidopsis thaliana. Nature 477, 419–423 (2011).

34. Kurtz, S. et al. Versatile and open software for comparing large genomes. Genome biology 5, R12 (2004).

35. Simão, F.A., Waterhouse, R.M., Ioannidis, P., Kriventseva, E.V. & Zdobnov, E.M. BUSCO: assessing genome assembly and annotation completeness with single-copy orthologs. Bioinformatics 31, 3210–3212 (2015).

36. Nattestad, M. & Schatz, M.C. Assemblytics: a web analytics tool for the detection of assembly-based variants. bioRxiv, 044925 (2016).

37. Li, R. et al. De novo assembly of human genomes with massively parallel short read sequencing. Genome research 20, 265–272 (2010).

38. Kajitani, R. et al. Efficient de novo assembly of highly heterozygous genomes from whole-genome shotgun short reads. Genome Res 24, 1384–1395 (2014).

39. Fondon, J.W.III, Martin, A., Richards, S., Gibbs, R.A. & Mittelman, D. Analysis of microsatellite variation in Drosophila melanogaster with population-scale genome sequencing. PLoS One 7, e33036 (2012).

40. Stanke, M. & Waack, S. Gene prediction with a hidden Markov model and a new intron submodel. Bioinformatics 19, ii215–ii225 (2003).

41. Dobin, A. et al. STAR: ultrafast universal RNA-seq aligner. Bioinformatics 29, 15–21 (2013).

42. Bowers, J.E. & Meredith, C.P. The parentage of a classic wine grape, Cabernet Sauvignon. Nature genetics 16, 84–87 (1997).

43. Jaillon, O. et al. The grapevine genome sequence suggests ancestral hexaploidization in major angiosperm phyla. Nature 449, 463–467 (2007).

44. Patel, S., Swaminathan, P., Fennell, A. & Zeng, E. in Bioinformatics and Biomedicine (BIBM), 2015 IEEE International Conference on 1771–1773 (IEEE, 2015).

45. Garrison, E. & Marth, G. Haplotype-based variant detection from short-read sequencing. arXiv preprint arXiv:1207.3907 (2012).

46. Bansal, V. & Bafna, V. HapCUT: an efficient and accurate algorithm for the haplotype assembly problem. Bioinformatics 24, i153–159 (2008).

47. Degner, J.F. et al. Effect of read-mapping biases on detecting allele-specific expression from RNA-sequencing data. Bioinformatics 25, 3207–3212 (2009).

48. Liu, Y.-G. & Whittier, R.F. Rapid preparation of megabase plant DNA from nuclei in agarose plugs and microbeads. Nucleic acids research 22, 2168 (1994).

49. Hayward, G.S. Unique double-stranded fragments of bacteriophage T5 DNA resulting from preferential shear-induced breakage at nicks. Proceedings of the National Academy of Sciences 71, 2108–2112 (1974).

50. Myers, G. in Algorithms in Bioinformatics 52–67 (Springer, 2014).

51. Chaisson, M.J. & Tesler, G. Mapping single molecule sequencing reads using basic local alignment with successive refinement (BLASR): application and theory. BMC bioinformatics 13, 238 (2012).

